# Basolateral amygdala astrocytes are engaged by the acquisition and expression of a contextual fear memory

**DOI:** 10.1101/2022.09.11.507456

**Authors:** Rebecca L. Suthard, Ryan A. Senne, Michelle D. Buzharsky, Angela Y. Pyo, Kaitlyn E. Dorst, Anh (Mia) H. Diep, Rebecca H. Cole, Steve Ramirez

**Affiliations:** Graduate Program for Neuroscience, Boston University, Boston, MA, 02215; Undergraduate Program in Neuroscience, Boston University, Boston, MA, 02215; Department of Biomedical Engineering, Boston University, Boston, MA, 02215; Department of Psychological and Brain Sciences; The Center for Systems Neuroscience; Neurophotonics Center, and Photonics Center, Boston University, Boston, MA, 02215

## Abstract

Astrocytes are key cellular regulators within the brain. The basolateral amygdala (BLA) is implicated in fear memory processing, yet most research has entirely focused on neuronal mechanisms, despite a significant body of work implicating astrocytes in learning and memory. In the present study, we used *in vivo* fiber photometry to record from amygdalar astrocytes across fear learning, recall, and three separate periods of extinction. We found that BLA astrocytes robustly responded to foot shock during acquisition, that their activity remained remarkably elevated across days in comparison to unshocked control animals, and that their increased activity persisted throughout extinction. Further, we found that astrocytes responded to the initiation and termination of freezing bouts during contextual fear conditioning and recall, and this behavior-locked pattern of activity did not persist throughout the extinction sessions. Importantly, astrocytes do not display these changes while exploring a novel context, suggesting that these observations are context or memory-dependent. Chemogenetic inhibition of fear ensembles in the BLA did not affect freezing behavior or astrocytic calcium dynamics. Overall, our work presents a real-time role for amygdalar astrocytes in fear processing and provides new insight into the emerging role of these cells in cognition and behavior.

**Significance Statement:** We show that basolateral amygdala astrocytes are robustly responsive to footshock, exhibit unique calcium event characteristics following contextual fear acquisition, and ramp up activity at the initiation and termination of freezing bouts during fear conditioning and recall. This astrocytic calcium response to freezing behavior is not observed during extinction sessions, despite unique calcium events continuing through three days of training compared to no-shock controls. We find that astrocytes display context specific changes in calcium signaling, but chemogenetic inhibition of BLA fear ensembles does not impact freezing behavior or calcium dynamics. These findings show that astrocytes play a key, real-time role in fear learning and memory.

## 1) Introduction

Memory acquisition, recall, and extinction are critical phases for information processing in the brain; disruption of any of these processes can lead to pathological states of cognition and behavior. Fear memories are one of the most well studied forms of memory, and have been shown to recruit numerous brain areas including the hippocampus and basolateral amygdala (BLA). Specifically, during Pavlovian fear conditioning, the CA1 and CA3 subregions of the hippocampus (HPC) relay information to the amygdala via the ventroangular pathway or through the entorhinal cortex (EC), which also projects to the prefrontal cortex (PFC). The amygdala then is thought to send output to the central amygdala (CeM) which in turn sends output to the lateral hypothalamus or periaqueducteal gray (PAG) to alter the sympathetic nervous system and gate freezing behavior, respectively. This complex fear circuitry is necessary for proper memory processing and these regions each differentially contribute to the behavioral expression of fear.

Recent work has demonstrated the active involvement of astrocytes in cognition and behavior by regulating synaptic plasticity, supporting metabolic homeostasis, modulating neurotransmitter action and releasing their own gliotransmitters to exert wide-ranging effects on the brain (Araque et. al., 2014; Bezzi et. al., 2001; Araque et. al., 2001; Araque et. al., 1999; Haydon et. al., 2001; Perea & Araque, 2005; Perea et. al., 2009; Parpura et. al., 1994; Porter et. al., 1997; Volterra et. al., 2005; Koizumi et. al., 2005; Covelo et. al., 2018; Di Castro et. al., 2011; Fellin et. al., 2004; Durkee et. al., 2019). Broadly, chemogenetic and optogenetic perturbations of astrocytic functioning have been shown to impair or enhance both recent and remote memory in the hippocampus, amygdala and prefrontal cortex (Kol et. al. 2020; Adamsky et. al., 2018; Li et. al. 2020; Martin-Fernandez et. al., 2017; Liao et. al., 2017; Fan et. al., 2021). The effects of these manipulations depend heavily on the brain region of interest, the signaling pathways perturbed and the time point of manipulation during behavior. Mounting evidence suggests that astrocytes may modulate local and projection-specific network activity in memory processes (Kol et. al., 2020; Martin-Fernandez et. al., 2017). For example, a recent study demonstrated that hippocampal dorsal CA1 (dCA1) astrocytic Gq activation during contextual fear conditioning is sufficient to promote long-term potentiation and enhance subsequent recall in mice, whereas neuronal activation does not (Adamsky et. al., 2018). Further research in the BLA has shown that fear conditioning itself downregulates astrocytic Rac1 to facilitate the formation of a conditioned fear memory (Liao et. al., 2017; Fan et. al., 2021). Additionally, BLA astrocytic Gq pathway activation during fear conditioning increased auditory memory, but not contextual memory retrieval, dissociating the role of these cells in multiple types of aversive learning (Lei et. al., 2022). Finally, Gq activation of astrocytes in the BLA after cued fear extinction training decreases freezing levels during extinction recall 24 hours later (Shelkar et. al., 2021). This evidence supports bidirectional astrocyte-neuron communication in multiple subdivisions of the amygdala and suggests their active control over the functional connectivity of the amygdala with other canonical fear-learning ‘hubs’.

Despite the recent interest in astrocytic contributions to memory, these studies predominantly use perturbation approaches (e.g. cell-type-specific activation or inhibition of cellular activity), whereas neuronal investigations may now multiplex these causal approaches with optical imaging to gain real-time insight on cellular activity during behavior. Still, there are relatively few studies utilizing these approaches, even with newer genetically-encoded calcium indicators (GECIs) and transgenic mouse lines that are capable of preferentially targeting astrocytes for dynamic calcium recordings across behavior (Corkrum et. al., 2020; Lin et. al., 2021; Qin et. al., 2020; Lines et. al., 2020; Tsunematsu et. al., 2021). Understanding the activity of astrocytes in real-time is essential for understanding cognition, especially given that the brain predominantly consists of glia.

The BLA is a key hub for valence-specific memories. Prior work has shown that the BLA is necessary for the encoding and retrieval of the emotional component of fearful experiences, and lesioning experiments have shown that its disruption strongly inhibits proper emotional responses (Zhang and Li, 2018; Maren, Ahranov, and Fanselow 1996; Maren 1999). Furthermore, the BLA has also been shown to be necessary for the acquisition and extinction of contextual fear memory in mice, suggesting it plays a key role in every stage of fear learning. Despite the relatively large body of literature implicating the BLA in fear conditioning, almost all of this work has focused on neuronal responses, and there is currently a limited understanding on how astrocytic calcium responses in the BLA manifest across fear acquisition, recall, and contextual extinction. Furthermore, previous work suggests that perturbation of BLA neurons during fear recall is capable of diminishing freezing behaviors (Han et al. 2009, Gore et al. 2015; Liu et. al., 2022), although coupled activity at the astrocyte-neuron interface remains relatively understudied activity.

To address this, we use freely-moving fiber photometry (Gunaydin et. al., 2014; Cui et. al., 2014) to record population-level astrocytic calcium dynamics across the classic contextual fear conditioning (CFC) paradigm. First, we find that astrocytes in the BLA are shock-responsive, which suggests that astrocytes in this amygdalar sub-region process salient and/or aversive-related stimuli. Next, we find that astrocytes in the shock condition displayed unique calcium events across fear learning compared to the unshocked control group. Then we observed calcium peri-events at the initiation and termination of freezing bouts during recall, but this did not persist into extinction sessions. We then utilized activity-dependent and chemogenetic-mediated inhibition of BLA cells to determine if this affected animal behavior and/or astrocytic signaling, and found no change in freezing or real-time dynamics. Finally, astrocytic changes were context-dependent in nature, as in a novel context we observed no differences in exploratory behaviors, nor any changes in calcium event characteristics across the shock and no-shock groups.

Together, our experiments provide a more comprehensive understanding of the contributions of glial cells to learning and memory processes. Perturbation of these cells during extinction memory formation and maintenance may pave the way for more successful therapeutic interventions in humans with disorders of maladaptive fear learning, such as Post-Traumatic Stress Disorder (PTSD).

## 2) Materials and Methods

### 2.1) Animals

Wild type, male C57BL/6J mice (P29-35; weight 17-19g; Charles River Laboratories) were housed in groups of 4-5 mice per cage. The animal facilities (vivarium and behavioral testing rooms) were maintained on a 12:12 hour light cycle (0700-1900). Mice received food and water *ad libitum* before and after surgery. Following surgery, mice were group-housed with littermates and allowed to recover for 3 weeks before experimentation. All subjects were treated in accord with protocol 201800579 approved by the Institutional Animal Care and Use Committee (IACUC) at Boston University.

### 2.2) Stereotaxic Surgery

For all surgeries, mice were anesthetized with 3.0% isoflurane inhalation during induction and maintained at 1-2% isoflurane inhalation through stereotaxic nose cone delivery (oxygen 1L/min). Ophthalmic ointment was applied to the eyes to provide adequate lubrication and prevent corneal desiccation. The hair on the scalp above the surgical site was removed using Veet hair removal cream and subsequently cleaned with alternating applications of betadine solution and 70% ethanol. 2.0% lidocaine hydrochloride (HCl) was injected subcutaneously as local analgesia prior to midsagittal incision of the scalp skin to expose the skull. 0.1mg/kg (5mg/mL) subcutaneous (SQ) dose of meloxicam and 0.1-0.2 mL of sterile saline were administered at the beginning of surgery. For fiber photometry implant surgeries, animals received a unilateral craniotomy with a 0.5-0.6 mm drill-bit for basolateral amygdala (BLA) injections. A 10μL airtight Hamilton syringe with an attached 33-gauge beveled needle was slowly lowered to the coordinates of BLA: −1.40 anteroposterior (AP), −3.20 mediolateral (ML) and −4.80 dorsoventral (DV). All coordinates are given relative to bregma (mm). A volume of 500nL of AAV-GfaABC1D-cyto-GCaMP6f-SV40 (Penn Vector Core) was injected at 50nL/min using a microinfusion pump for the BLA coordinate (UMP3; World Precision Instruments). After the injection was complete, the needle remained at the target site for 5-7 minutes post-injection before removal. Following viral injection, a unilateral optic fiber (200μm core diameter; 1.25mm ferrule diameter) was implanted at the site of injection. The implant was secured to the skull with a layer of adhesive cement (C&M Metabond) followed by multiple layers of dental cement (Stoelting). Following surgery, mice were injected with a 0.1mg/kg intraperitoneal (IP) dose of buprenorphine for pain management. They were placed in a recovery cage with a heating pad on medium heat until fully recovered from anesthesia. To allow for recovery and viral expression, we waited 3 weeks before beginning our behavioral paradigm. Histological assessment verified viral targeting and data from off-target injections were not included in further analyses.

For activity-dependent (‘engram’) virus surgeries, mice were placed on doxycycline (Dox) diet for 72 hours prior to surgery. The same injection, viral volume and fiber photometry implantation methods were used as described above, with the addition of 200-250nL AAV9-c-fos-tTA-TRE-hM4Di-mCherry or AAV9-c-fos-tTA-TRE-mCherry virus in bilateral BLA to allow for inhibition of the fear engram during recall.

### 2.3) Fiber Photometry

A 470-nm LED (Neurophotometrics; FP3002) delivered an excitation wavelength of light to astrocytes expressing GCaMP6f via a single fiber optic implant. The emitted 530-nm signal from the indicator was collected via this same fiber, spectrally-separated using a dichroic mirror, passed through a series of filters and was focused on a scientific camera. Calcium-independent isosbestic signals were simultaneously captured by alternating excitation with 415-nm LED to dissociate motion, tissue autofluorescence, and photobleaching from true changes in fluorescence. All wavelengths were interleaved and collected simultaneously using Bonsai (Lopes et. al., 2015). The sampling rate for the signals was 28Hz (28 frames per second). Time series were analyzed using an in-house pipeline and fluorescence signals were normalized to the median for comparison of event amplitude (peak height; % dF/F), frequency (Hz), total fluorescence (area under the curve; AUC) and duration (full-width half maximum; FWHM). Statistical analyses were performed using Python and data reported as Mean ± SEM.

### 2.4) Behavioral Testing

#### Fear Conditioning, Recall, Extinction Experiments

On Day 1, mice were placed into the shock context (Cxt A) where they underwent a 360s contextual fear conditioning session. Footshocks (1.5mA, 2s duration) were administered at the 120, 180, 240 and 300 second time points at 1.5mA intensity. On Day 2, mice were placed back in Cxt A for 360s of recall where they received shock on the previous day. There were no shocks administered during this session. On Days 3-5, mice were placed back in Cxt A for 900s without shock administration to extinguish the fear memory across days. At the completion of extinction testing, mice in the no-shock group were administered a single 1.5mA footshock to confirm the presence of calcium signal. On Day 6, mice underwent a 600s session of exploration in a novel open field context (Cxt B) to assess the context-dependent nature of astrocytic calcium changes, if present.

Days 1-5 took place in mouse conditioning chambers (18.5 x 18.5 x 21.5cm)(Coulbourn Instruments) with metal-panel side walls, plexiglass front and rear walls and a stainless-steel grid floor composed of 16 grid bars that were connected to a precision animal shocker that delivered the four foot shocks. A video camera was mounted on a tripod in a front-facing orientation to the conditioning chamber for fear conditioning, recall and extinction. Day 6 took place in an open field arena (61cm x 61cm) with black plastic walls and a taped area (45cm x 45cm) in the middle delineating a “center” region. For this session, a top-down camera was used to record behavioral video during the session. The chambers were cleaned with 70% ethanol solution prior to each animal placement. All behavioral testing was performed during the animal’s light-cycle.

#### Engram behavioral experiments

Mice were separated into two groups: hM4Di (experimental) and mCherry (control). On Day 1, mice were taken off of their Dox diet for 48 hours to open the ‘tagging window’. On Day 3, mice were placed in the same contextual fear conditioning (CFC) chamber described above, where they received four, 1.5mA foot shocks for 360 seconds. All mice were placed immediately back on Dox after this session. On Day 4, mice were placed back in the same conditioned context for a 360 second session of recall. Both groups of mice received a single I.P. injection (3 mg/kg) of clozapine-N-oxide (CNO) 30 minutes prior to the session to inhibit the ‘tagged’ fearful experience. 90 minutes after the start of recall, mice were transcardially perfused to capture peak cFos protein levels in the BLA due to inhibition of the tagged neuronal ensemble. Mice had calcium dynamics recorded across all three sessions using fiber photometry. For these sessions, a front-facing video camera was used to obtain behavioral videos during fear conditioning and recall, and a top-down facing video camera was used to obtain home cage video.

#### Chemogenetic Parameters

For chemogenetic silencing of bilateral BLA fear engram neurons, we used the inhibitory Designer Receptor Exclusively Activated by Designer Drugs (DREADDs), hM4Di. hM4Di drives inhibition of infected neurons when bound to the ligand, CNO (Sigma-Aldrich). A 0.6 mg/mL solution of CNO was prepared in sterile saline and 0.5% dimethyl sulfoxide (DMSO). All mice were I.P. injected with sterile saline for five days prior to experimentation for habituation. On the day of contextual recall, mice were injected with a 3 mg/kg dose of CNO solution 30 minutes prior to the start of the session to capture peak drug concentration.

### 2.5) Immunohistochemistry

On Day 6, mice were overdosed with 3% isoflurane and perfused transcardially with cold (4°C) phosphate-buffered saline (PBS) followed by 4% paraformaldehyde (PFA; pH = 7.4) in PBS. Brains were extracted and kept in PFA at 4°C for 24-48 hours and transferred to a 30% sucrose in PBS solution. Brains were sectioned into 50μm thick coronal sections with a vibratome and collected in cold PBS or 0.01% sodium azide in PBS for long-term storage. Sections were washed three times for 10-15 minutes with PBS to remove 0.01% sodium azide used for storage. Vibratome sections were incubated for 2 hours in PBS combined with 0.2% Triton (PBST) and 5% bovine serum albumin (BSA) on a shaker at room temperature. Sections were incubated in the primary antibodies (1:1000 mouse monoclonal anti-GFAP [NeuroMab]; 1:1000 rabbit polyclonal anti-Iba1 [Wako]; 1:500 guinea pig anti-NeuN [SySy]) diluted in PBST/1% BSA at 4°C for 24 or 48 hours. The slices were washed three times for 10-15 minutes each in 1xPBS. The secondary antibodies were diluted in secondary antibody solution (PBST/1% BSA) and incubated for 2 hours at room temperature. The following secondary antibodies were used: 1:1000 Alexa 555 anti-mouse [Invitrogen], 1:1000 Alexa 555 anti-rabbit [Invitrogen], 1:200 Alexa 555 anti-guinea pig [Invitrogen]. The sections were then washed three times with 1xPBS or PBST for 10-15 minutes each and mounted using Vectashield Mounting Medium with DAPI (Vector Laboratories). Once dry, slides were sealed with clear nail polish on each edge and stored in a slide box in the fridge (4°C). Mounted slices were imaged using a confocal microscope (Zeiss LSM800, Germany).

For activity-dependent (‘engram’) experiments, immunohistochemistry was performed as described above with the use of the following primary (1:1000 chicken anti-GFP [Invitrogen], 1:1000 guinea pig anti-RFP [SySy]) and secondary (1:200 Alexa 488 goat anti-chicken [Invitrogen], 1:200 Alexa 555 goat anti-guinea pig [Invitrogen]) antibodies to confirm co-expression of hM4Di-mCherry (RFP) and GfaABC1D-GCaMP6f (GFP) in the BLA.

Brains from all mice used in fiber photometry experiments were analyzed to check adequate fiber location and proper and selective viral expression. Animals that did not meet the criteria for proper fiber location and virus expression were discarded.

### 2.6) Imaging and Cell Counting

Imaging was performed using a Zeiss LSM 800 epifluorescence microscope with 10x and 20x objectives using the Len2.3 software. Full slice images of BLA GfaABC1D-GCaMP6f (green) and DAPI (Figure 1C) were captured with a 10x objective (6554 x 9319 pixels) along with a zoomed-in (1024 x 1024 pixels) single-tile z-stack at 20x magnification (Figure 1B).

**Figure 1.**
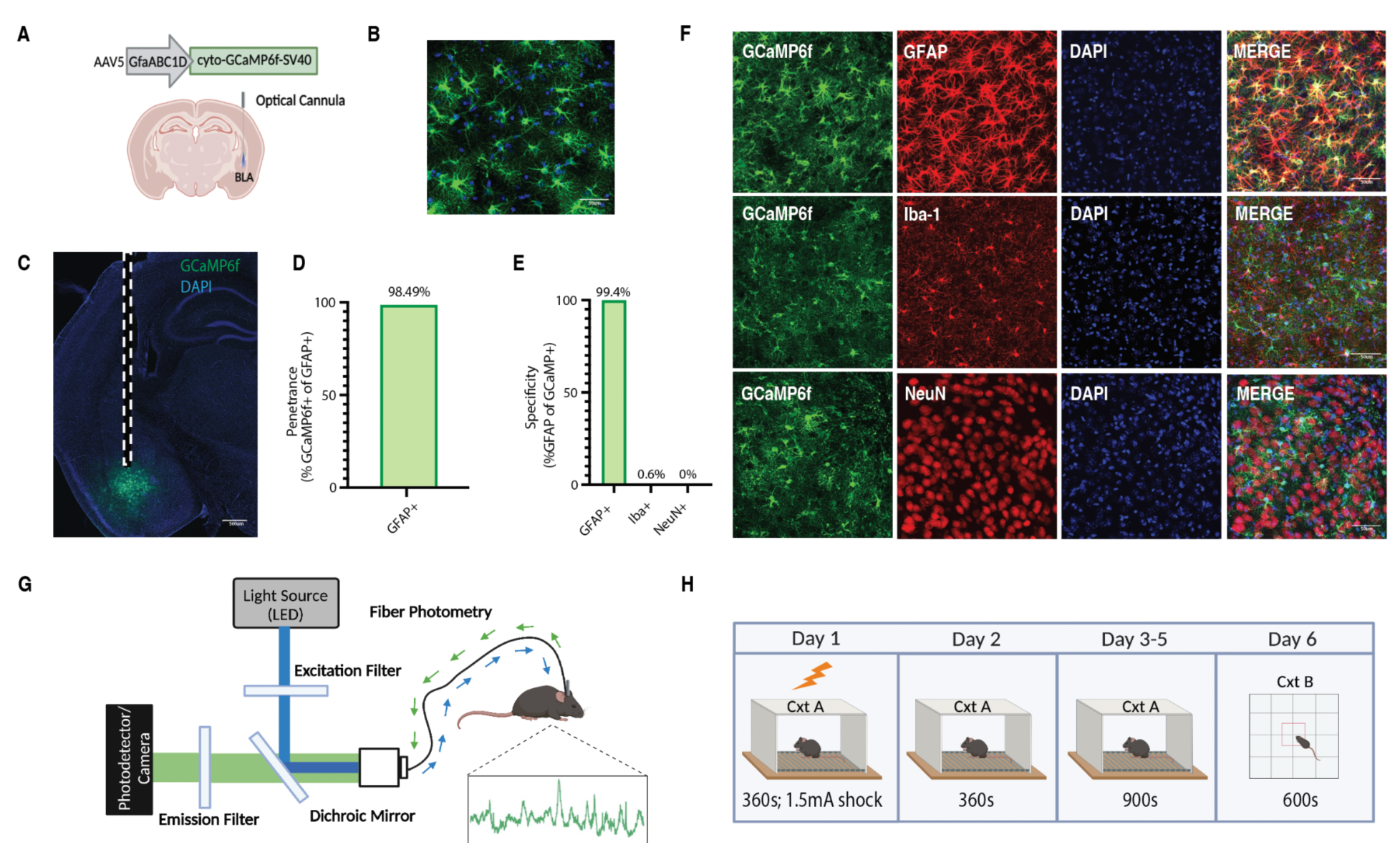
Population-level calcium recordings of basolateral amygdala astrocytes across contextual fear conditioning, recall and extinction. (A) Viral strategy and fiber implantation strategy for shock and no shock conditions. The genetically-encoded calcium indicator (GECI) AAV5-GfaABC1D-cyto-GCaMP6f-SV40 was unilaterally injected into the BLA region of wild type mice. (B) Representative image of GFAP-GCaMP6f+ (green) and DAPI+ (blue) cell expression within the BLA at 20x magnification and (C) 10x magnification; scale bar indicates 500 micrometers (um). Dashed white lines indicate the approximate location of the unilateral fiber implantation. (D) Penetrance of GCaMP6f (2251 GCaMP6f+/2381 GFAP+ = 98.49%) (n=3; 4 slices/mouse). (E) Specificity of GCaMP6f (4 Iba-1+/662 GCaMP6f+ = 0.604% microglia; 0 NeuN+/1064 GCaMP6f+ = 0.00% neurons; 2242 GFAP+/2256 GCaMP6f+ = 99.4% astrocyte)(n=3; 4 slices/mouse). (F) Representative expression of GCaMP6f expression, and overlap with microglial (Iba-1), astrocytic (GFAP) and neuronal (NeuN) markers; scale bar indicates 50 micrometers (um).(G) *In vivo* fiber photometry set-up; a 470-nm LED delivered an excitation wavelength to GCaMP6f-expressing astrocytes via a patch cord and single fiber optic implant in freely moving mice. The emitted 530-nm signal from the indicator was collected via the same patch cord and fiber, spectrally-separated using a dichroic mirror, passed through a series of filters and focused on a scientific camera. A representative calcium time series trace is shown for astrocytic calcium. Calcium-independent isosbestic signal was recorded simultaneously to account for motion, tissue autofluorescence and photobleaching across time. (H) Behavioral paradigm; mice underwent contextual fear conditioning (CFC) on Day 1 in Context A (Cxt A) for 360 seconds where they received 4, 1.5mA foot shocks. Day 2, mice were placed back into Cxt A for contextual recall for 360 seconds in the absence of foot shock. Days 3-5, mice underwent three contextual extinction sessions for 900 seconds each. Day 6, mice were placed into a novel open field context B (Cxt B) for 600 seconds. Mice were perfused and brains extracted for histological assessment.

The cellular specificity and penetrance of the GCaMP6f viral vectors (Figure 1D-F) was tested by immunohistochemical analysis of the BLA. These images of NeuN, Iba-1 and GFAP were captured in z-stacks of the BLA at 20x magnification. 20x magnification, single-tile (1024 x 1024 pixels) representative images are shown (Figure 1B). Cell counts for NeuN, Iba-1, GFAP and overlaps were performed using Ilastik, a machine-learning-based image analysis tool (Berg et. al., 2019). Of the 2256 cells expressing GCaMP6f (n = 3 mice; 3 slices), 2242 were astrocytes (identified by GFAP), resulting in a specificity of 99.4%. Of the 662 cells expressing GCaMP6f (n=3 mice; 3 slices), 4 were microglia (identified by Iba-1), resulting in a specificity of 0.6%. Of the 1064 cells expressing GCaMP6f (n = 3 mice; 3 slices), 0 were neurons (identified by NeuN), resulting in a specificity of 0%. Finally, of 2381 GFAP+ astrocytes, 2251 were GCaMP6f+ (n=3 mice; 3 slices), resulting in a penetrance of 98.49%.

For engram experiments, bilateral BLA was imaged at 20x magnification to confirm expression of hM4Di and mCherry and single-tile (1024 x 1024 pixels) images were shown (Figure 7B).

### 2.7) Behavioral Analysis

An automated video tracking system, AnyMaze, was used for supervised analysis of freezing bout initiation and termination in the shock context (Cxt A), as well as the total distance traveled, mean speed, number of entries into the center and total time spent in the center of the open field (Cxt B). Additionally, the pose estimation algorithm, DeepLabCut, was used to perform unbiased animal behavioral evaluation of kinematics (position, acceleration, velocity) for use in generalized linear modeling (GLM) (Mathis et. al., 2018). This behavioral data was time locked to our fiber photometry time series data for all analyses.

### 2.8) Statistical and Fiber Photometry Analysis

All statistical analysis and subsequent photometry analysis was performed via custom python scripts freely available at https://github.com/rsenne/RamiPho. Our environment was built off of scipy (ver. 1.7.3), statsmodels (ver. 0.13.2), numpy (ver. 1.22.3), pandas (ver. 1.4.2), matplotlib (ver. 3.5.1), numba (ver. 0.55.2), and seaborn (ver. 0.11.2).

All photometry signals were baseline corrected using an adaptive iteratively reweighted penalized least squares method (Zhang, Chen, and Liang, 2010). We then performed a simple kernel smoothing to increase the signal to noise ratio. For “event detection” we used a method published in prior literature (Howe et al. 2019). DF/F was calculated by subtracting the median of the trace from the current fluorescence value and then dividing by the median, giving a percent difference from the median or “baseline”. We then found any peaks greater than 1 standard deviation away from the mean, in both the positive and negative direction. We then determined the minimum width such that the ratio of positive to total transients (negative and positive) was greater than or equal to 0.99. Any transients below this width were subsequently discarded.

For peri-event analysis, we used a tCI method as proposed previously (Jean-Richard-dit-Bressel, Clifford & McNalley, 2020). For each window of interest, we calculated a 95%, 99% and 99.9% confidence interval. If the CI did not contain the null assumption (dF/F = median of the event window), for a period greater than 0.5 s we concluded that a significant peri-event occurred. This type of method does not allow an exact p-value and thus is omitted from the text.

For generalized linear modeling (GLM), we fit a model using a Gaussian family and identity link function. We hypothesized that the calcium activity (dF\F) could be explained by a combination of the isosbestic signal, velocity and acceleration (up to the third degree), the binary freezing values, the animal identification number, and an interaction effect between the animal and a series of basis spline functions for: the 10 seconds following shock presentation, and 5 seconds following initiation of freezing behavior, and 5 seconds following the termination of freezing. We chose to fit a singular model as opposed to individuals models for each model because we wanted to maximize the number of samples to get better fits for coefficients that should be shared across animals (e.g. the contribution of velocity to the trace) while still including the animal specific terms to account for individual variability within the SNR of each trace (e.g. the interaction effect between the animals and splines will give unique coefficients for each animal helping to model discrete events better). <u>Thus, our model can be summarized as follows</u>:

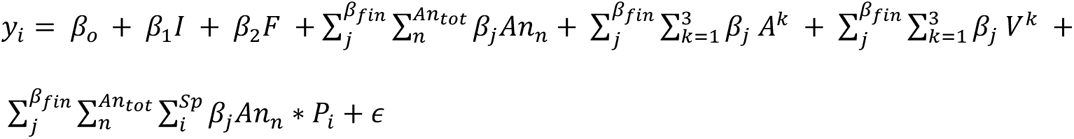

Where I is the isosbestic channel, F is the freezing (binary: 1 is freezing, else 0), An is a dummy variable encoding animal ID, A is acceleration, V is velocity, P is our polynomial spline functions, k degree of the polynomial function applied to the kinematic variables, An_tot is the total number of animals, Sp is the total number of splines, and \beta_fin is the final index for the specific summation. Or more simply stated: dF/F ∼ Isosbestic + Freezing + Animal + Acceleration Functions + Velocity Functions + Animal Specific Splines + Error.

The above model is now considered the “full model” such that this is the highest order model possible for our analysis. To do model selection, simpler models were tested, and when nested, a maximum likelihood ratio test was performed (MLRT) to determine the final model. If comparing two models that were not nested, the Akaike Information Criterion (AIC) was used.

## 3) Results

Astrocytes in the basolateral amygdala are actively involved in fear memory (Liao et. al., 2017; Fan et. al., 2021; Lei et. al., 2022; Shelkar et. al., 2021; Stehberg et. al., 2012), but their real-time dynamics during the acquisition and maintenance of conditioned fear in mice is relatively unknown. We monitored astrocyte calcium levels in the BLA using fiber-photometry in freely behaving mice across the acquisition, retrieval, and extinction of conditioned fear. Wild type mice were injected unilaterally with AAV-GfaABC1D-cyto-GCaMP6f to express the genetically-encoded calcium indicator (GECI) GCaMP6f selectively in astrocytes (Figure 1A-C). To quantify the penetrance and specificity of our viral system, we co-labeled GCaMP6f+ cells with GFAP, Iba-1 and NeuN, markers for astrocytes, microglia and neurons, respectively (Figure 1D-F). Of 2381 GFAP+ cells, 2251 were GCaMP6f+ (n=3; 4 slices/mouse), resulting in a penetrance of 98.49% (Figure 1D, F). Of 662 GCaMP6f+ cells, 4 were Iba1+, resulting in a specificity of 0.604% for microglia (Figure 1E-F). Of 1064 GCaMP+ cells, 0 were NeuN+, resulting in a specificity of 0.0% for neurons (Figure 1E-F). Finally, of 2256 GCaMP+ cells, 2242 were GFAP+, resulting in a specificity of 99.4% for astrocytes (Figure 1E-F).

To test the hypothesis that astrocytes play an active role in the acquisition and maintenance of contextual fear, we used *in vivo* fiber photometry to record their activity across all experimental days of our behavioral task (Figure 1G-H). Recent literature using calcium imaging in ventral hippocampus (vHPC) has shown that a subset of BLA-vHPC projecting neurons were responsive to aversive shock during CFC (Jimenez et. al., 2020). This provides ample evidence that BLA neurons and astrocytes are also likely to be shock-responsive. On Day 1, mice underwent contextual fear conditioning with the administration of 1.5mA foot shocks at 120, 180, 240 and 300 second timepoints (Figure 1H). To assess whether BLA astrocytes responded specifically to footshock, we performed peri-event analysis to determine if calcium transients were temporally locked to footshock. We used a tCI confidence interval method to classify significant perievents around a time-point of interest as previously described (Jean-Richard-dit-Bressel, Clifford & McNalley, 2020; *See Methods*). Astrocytes in the shock group displayed robust increases in population-level calcium at the onset of each foot shock during the session compared to the no-shock condition, as shown by a representative calcium time series (% dF/F) from each group (Figure 2A) and raster plots (z-scored % dF/F) including all mice from each group (Figure 2D-E). Further analysis revealed that the shock group had significantly increased % dF/F from baseline after the onset of each foot shock compared to the no-shock condition (Footshock peri-event analysis; 99% confidence interval (CI))(Figure 2B). Specifically, the shock group had an increase of 44.6% dF/F from baseline after the onset of each footshock, on average, compared to 1.6% increase dF/F for the no-shock condition (Independent samples t-test; Welch’s correction, p=0.0022) (Figure 2C).

**Figure 2.**
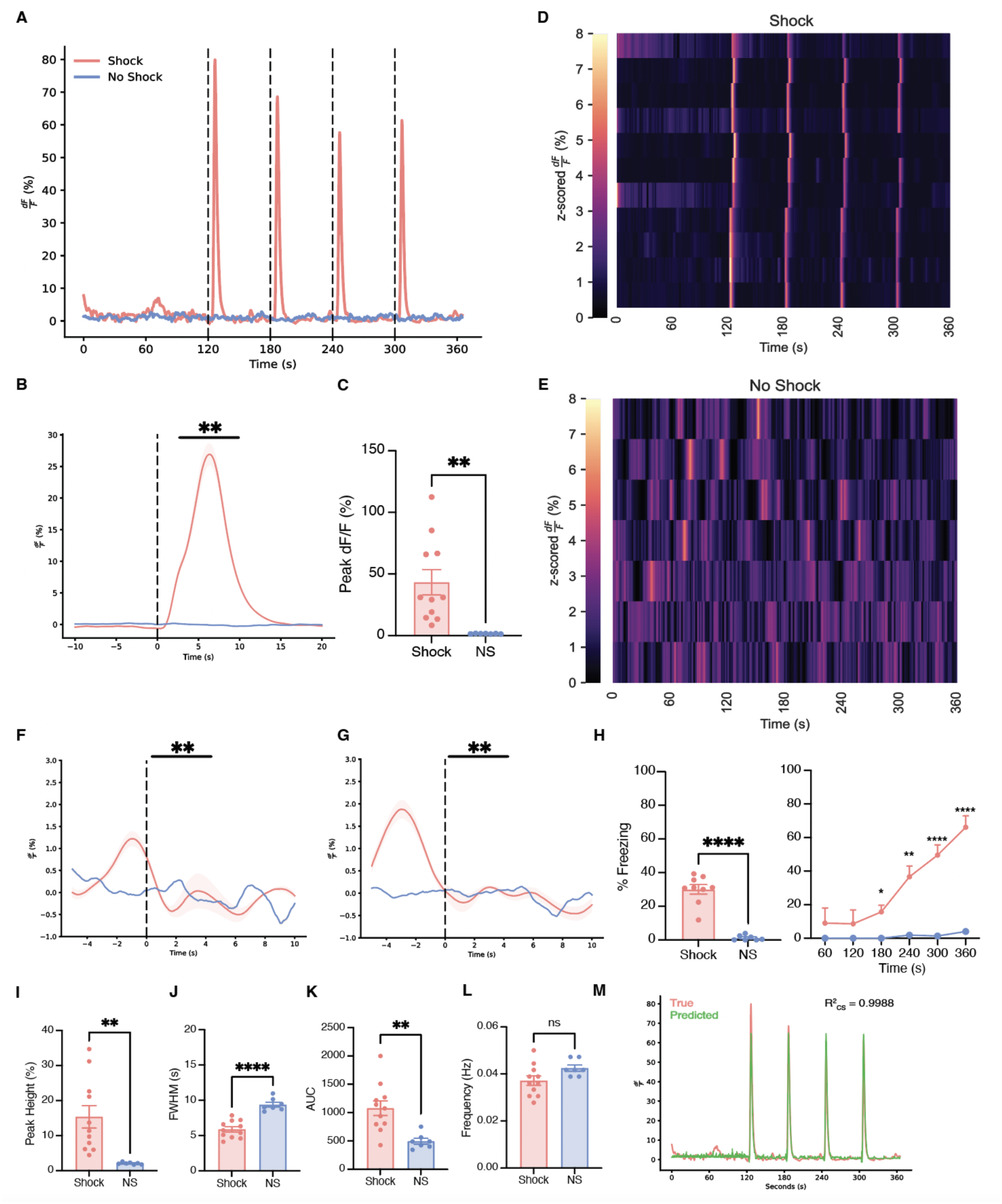
Basolateral amygdala astrocytes robustly respond to foot shock during contextual fear conditioning and exhibit unique calcium event dynamics compared to no-shock controls. (A) Representative calcium time series (dF/F %) for shock and no-shock conditions during the 360 second CFC session. 1.5mA foot shocks occurred at the 120, 180, 240 and 300 second time points, as indicated by vertical dashed lines. (B) Peri-event analysis for 1.5 mA foot shock, with the onset of foot shock occurring at the dashed line (time = 0). (C) Quantification of the average percent change in peak dF/F at the onset of foot shock. (D-E) Z-scored dF/F (%) across CFC for (D) shock and (E) no-shock conditions; each row represents a single subject across time within the session. (F-G) Peri-event analysis for the initiation (C) and termination (E) of freezing behavior, with each event occurring at the dashed line (time = 0). (H) Average percent freezing (left) and freezing across time within the CFC session (right). (I-L) Calcium event metrics; (I) peak height, (J) full-width half maximum, (K) area under the curve, and (L) frequency. (M) True and predicted traces produced from a generalized linear model. All error bars and bands indicate SEM. For t-tests and ANOVAs, p ≤ 0.05, **p ≤ 0.01, ***p ≤ 0.001, ****p ≤ 0.0001, ns = not significant. For peri-event metrics, * = 95% CI; ** = 99% CI; ns = not significant. For t-tests and ANOVAs, shock n=11, no-shock n=7. For foot shock peri-events, shock n=11, no-shock n=7. For freezing peri-events, shock n=11; no-shock n=2 due to mice not freezing during this session.

We next tested the hypothesis that astrocytes modulate their behavior in response to the initiation and/or termination of freezing. There were significant peri-events, where the event started prior to the initiation of freezing (Freeze initiation peri-event analysis; 99% CI)(Figure 2F) and the termination of freezing (Freeze termination peri-event analysis; 99% CI), which indicates that astrocytic signaling becomes coupled to freezing behavior during fear memory acquisition (Figure 2G). Behaviorally, mice in the shock group had a higher freezing across the 360s session, compared to the no-shock group that did not experience a foot shock (Independent t-test; p<0.0001)(Figure 2H). The shock group successfully acquired fear across the CFC session compared to the no-shock condition (Two-way ANOVA with repeated measures (RM); Interaction: F (5, 80) = 10.14, p<0.0001; Time bin: F (1.750, 28.00) = 13.02, p=0.0002; Group: F (1, 16) = 23.59, p=0.0002; Subject: F (16, 80) = 4.593, p<0.0001)(Figure 2H). *Post-hoc* analysis demonstrated significant group differences in freezing at the 180, 240, 300 and 360 second time bins (Sidak’s multiple comparisons: 180s: p=0.0167; 240s: p=0.0015; 300s: p<0.0001; 360s: p<0.0001)(Figure 2H).

We calculated event metrics such as peak height, full-width half max (FWHM), area under the curve (AUC), and frequency (events/minute) for shock and no-shock groups. The shock group had significantly increased event peak height, increased AUC, and decreased full-width half max (FWHM) compared to no shock during CFC ([Peak height: Independent t-test; Welch’s correction, p=0.0019][AUC: Independent t-test; Welch’s correction, p=0.0011][FWHM: Independent t-test; p<0.0001][Frequency: Independent t-test; p=0.069 (ns)](Figure 2I-L).To further supplement these analyses, we tested if the astrocytic calcium signals could be decoded as a linear combination of the isosbestic channel, freezing, animal identification number (ID), kinematics, and a set of polynomial spline functions to model the shock response, and the initiation and termination of freezing bouts. For a full elaboration of the model and model selection, see *Methods Section 2.8*. We found that during fear conditioning, the most parsimonious model that could explain the most variability in the dataset contained only the animal specific splines and kinematic information up to the third degree, meaning that these signals almost entirely encode the shock information and some amount of motion related output. (R^2^_CS_=0.9988)(Figure 2M). Together, the calcium events for the shock condition (i.e. higher amplitude, higher total fluorescence, shorter duration events) suggest that astrocytes become more active after the presentation of a salient stimulus, as suggested by previous literature demonstrating that these cells in dCA1 are responding in a stimulus-dependent manner (Adamsky et. al., 2018).

Recent studies have shown that manipulation of astrocytes during retrieval of a conditioned fear memory does not affect recent or remote memory recall, though their activity within a given session remained unmeasured (Adamsky et. al., 2018; Kol et. al., 2020). To that end, we investigated the real-time dynamics of these cells in mice that received foot shock vs. neutral exposure to the same context. On Day 2 of our experiment, mice underwent contextual recall without the presence of the unconditioned stimulus (US; foot shock)(Figure 1H). When comparing population-level calcium activity across the shock and no-shock groups, we observed stark differences in the engagement of astrocytes (Figure 3A-B, D). Specifically, we observed that the presence of the US (i.e. footshock) during fear conditioning continued to engage astrocytes when placed back in the original context, while the no-shock group displayed low levels of calcium activity. As the US is the most salient manipulated variable between these two groups, we speculate that astrocytes may be engaged in a learning-dependent manner within the basolateral amygdala (BLA). We also tested the hypothesis that astrocytes modulate their behavior in response to the initiation and/or termination of freezing. There were significant peri-events, where the event started prior to the initiation of freezing (Freeze initiation peri-event analysis; 99% CI)(Figure 3C) and immediately after the initiation of freezing (Freeze termination peri-event analysis; 99% CI)(Figure 3E). Behaviorally, mice in the shock group exhibited increased freezing compared to the no-shock condition that did not receive the CS-US pairing (Mann-Whitney U-test; p<0.0001)(Figure 3F). Across the recall session, mice in the shock group maintained higher levels of freezing compared to no-shock controls (Two-way ANOVA RM; Interaction: F (5, 80) = 1.163, p=0.3346; Time Bin: F (5, 80) = 1.906, p=0.1024; Group: F (1, 16) = 93.18, p<0.0001; Subject: F (16, 80) = 8.924, p<0.0001) (Figure 3G). *Post-hoc* analysis across groups supported significantly higher levels of freezing in the shock condition during recall across all time bins (Sidak’s multiple comparisons: 60s: p<0.0001; 120s: p<0.0001; 180s: p<0.0001; 240s: p<0.0001; 300s: p<0.0001; 360s: p<0.0001) (Figure 3G). Furthermore, to quantify these differences in the traces across groups, we calculated the same average event metrics mentioned above (i.e. peak height, FWHM, AUC, frequency). The shock group had increased average peak height, decreased FWHM, increased AUC, and there were no differences in frequency of events across the session ([Peak height: Independent t-test; Welch’s correction, p=0.0009][AUC: Independent t-test; Welch’s correction, p=0.0156][FWHM: Independent t-test; Welch’s correction, p=0.0005][Frequency: Independent t-test; Welch’s correction, p=0.3642).(Figure 3H-K). This suggests that after context-shock association, astrocytes retain similar calcium dynamics on the following day when placed back in the conditioned environment in the absence of shock. We then fit a Gaussian generalized linear model (GLM) as previously described, to determine which variables best accounted for our calcium signal. Interestingly we saw that the kinematic information, freezing, freezing initiation and termination splines, and animal information best accounted for the recall data (R^2^_CS_=0.3059)(Figure 3L). This indicates that during recall there does not seem to be any temporal encoding of the shock that we can detect with this analysis, and further suggests that the activity of these astrocytes is tethered to freezing bouts.

**Figure 3.**
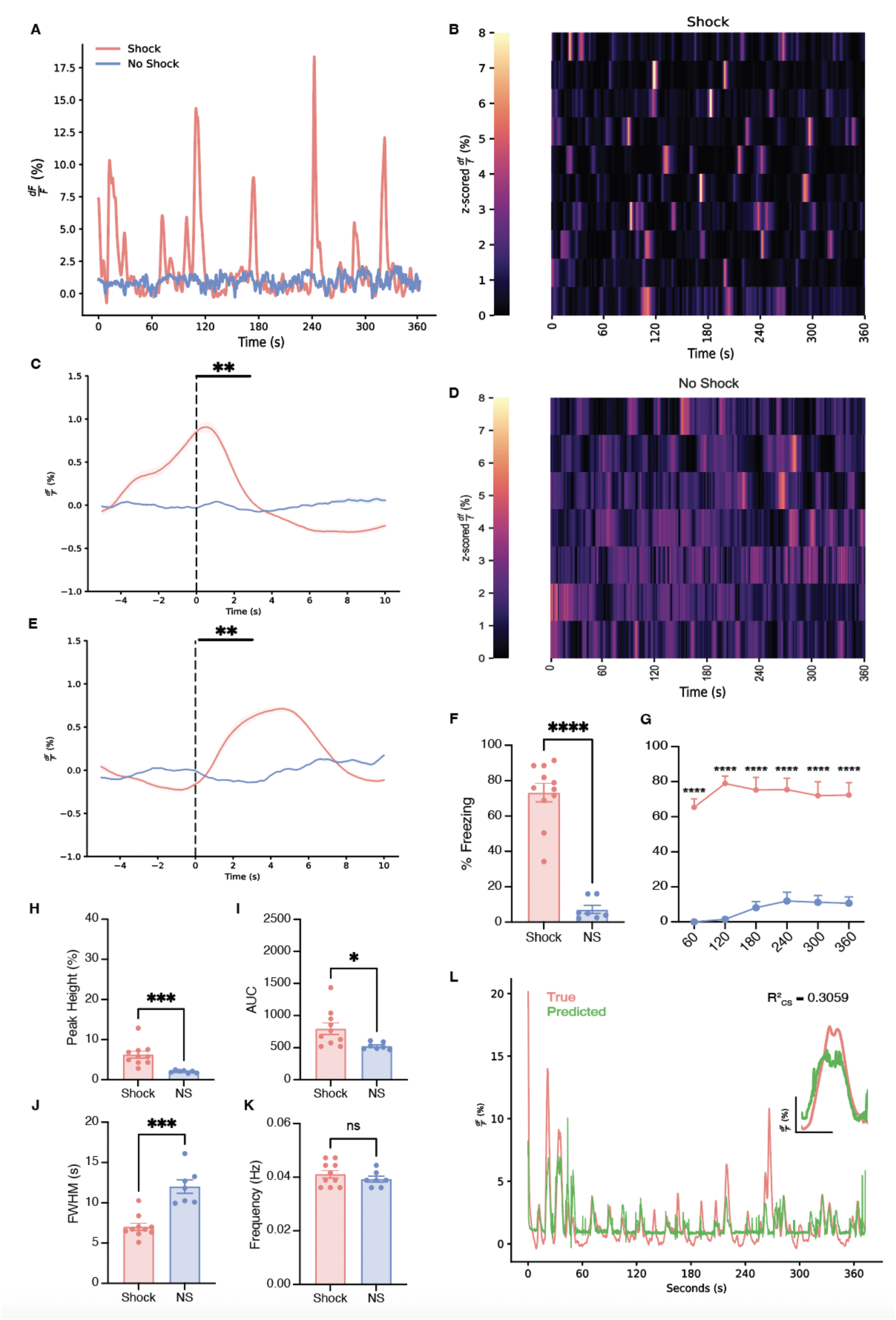
BLA astrocytes respond reliably to the initiation and termination of freezing behavior during contextual recall. (A) Representative calcium time series (dF/F %) for shock and no-shock conditions during the 360 second recall session in the absence of foot shock. (B, D) Z-scored dF/F (%) across recall for (B) shock and (D) no-shock conditions; each row represents a single subject across time within the session. (C, E) Peri-event analysis for the initiation (C) and termination (E) of freezing behavior, with each event occurring at the dashed line (time = 0). (F-G) Behavioral analysis; (F) average percent freezing and (G) freezing across time within the recall session. (H-K) Calcium event metrics; (H) peak height, (I) area under the curve, (J) full-width half maximum, and (K) frequency. (L) True and predicted traces produced from a generalized linear model. Error bars indicate SEM. For t-tests and ANOVAs, p ≤ 0.05, **p ≤ 0.01, ***p ≤ 0.001, ****p ≤ 0.0001, ns = not significant. For peri-event metrics, * = 95% CI; ** = 99% CI;, ns = not significant. For t-tests and ANOVAs, shock n=11, no-shock n=7. For freezing peri-events, shock n=11, no-shock n=6 due to a mouse not freezing during the recall session.

To further evaluate the experience-dependent role of astrocytic calcium across the extinction of contextual fear, mice underwent three days of extinction on Days 3-5 of our behavioral paradigm (Figure 1H). When comparing astrocytic calcium levels during extinction, we observed higher population activity in the shock group in the absence of the original US compared to the no-shock condition across all three days (Figure 4A-I). This suggests that astrocytes are continuing to be engaged in the shock group, and perhaps in a memory-dependent manner as an extinction memory is being formed across days. The no shock group displayed minimal calcium activity, which may be due to continued exploration of a novel environment as it becomes familiar (Qin et. al., 2020). Furthermore, to quantify these differences across groups, we calculated event metrics for each extinction session. For extinction day 1, the shock condition had calcium events with increased peak height, increased AUC, decreased FWHM (i.e. duration of event), and decreased frequency ([Peak height: Independent t-test; Welch’s correction, p=0.0016][AUC: Independent t-test; Welch’s correction, p=0.0036][FWHM: Mann-Whitney U-test, p=0.0019][Frequency: Independent t-test, p=0.0253])(Figure 4J-M). For extinction day 2, the shock condition had calcium events with increased peak height, increased AUC, decreased FWHM and no difference in frequency ([Peak height: Independent t-test, p=0.0001][AUC: Independent t-test, p<0.0001][FWHM: Mann-Whitney U-test, p=0.0043][Frequency: Independent t-test, p=0.2050])(Figure 4J-M). Finally, for extinction day 3, the shock condition had calcium events with increased peak height, but no significant differences in FWHM, AUC or frequency compared to no-shock controls ([Peak height: Independent t-test, p=0.0010][AUC: Independent t-test; Welch’s correction, p=0.126][FWHM: Independent t-test, p=0.1447][Frequency: Independent t-test, p=0.3851)(Figure 4J-M). Interestingly, this suggests that astrocytes are initially impacted by the presence of the foot shock during CFC, but do not adapt further across extinction days. To further investigate astrocytic calcium responses to freezing behaviors, peri-event metrics for the initiation and termination of freezing were calculated as previously performed in contextual recall. Interestingly, astrocytic calcium in the shock group was not responsive to the initiation or termination of freezing behavior for all extinction sessions (Freeze initiation and termination peri-event analysis; ns)(Figure 5A-B, D-E, G-H). Behaviorally, mice in the shock group exhibited higher average levels of freezing than the no-shock condition group for all three days ([Extinction 1: Independent t-test; Welch’s correction, p<0.0001][Extinction 2: Independent t-test; Welch’s correction, p=0.0042][Extinction 3: Independent t-test; Welch’s correction, p=0.0123])(Figure 5J). For each extinction session, shocked mice displayed higher levels of freezing at each time bin than no-shock controls ([Extinction 1: Two-way ANOVA RM; Interaction: F (14, 210) = 1.140, p=0.3247; Time bin: F (4.530, 67.95) = 1.726, p=0.1467; Group: F (1, 15) = 22.92, p=0.0002; Subject: F(15, 210) = 18.83, p<0.0001)][Extinction 2: Two-way ANOVA RM; Interaction: F (14, 168) = 1.204, p=0.2765; Time Bin: F (4.306, 51.68) = 1.364, p=0.2574; Group: F (1, 12) = 18.86, p=0.0010; Subject: F (12, 168) = 30.92, p<0.0001][Extinction 3: Interaction: F (14, 168) = 0.9191, p=0.5395; Time bin: F (3.186, 38.23) = 1.160, p=0.3392; Group: F (1, 12) = 12.14, p=0.0045; Subject: F (12, 168) = 25.16, p<0.0001])(Figure 5C, F, I). Specifically, *post-hoc* analysis for extinction days revealed specific time bin differences across groups, and this declines across extinction days ([Extinction 1: Sidak’s multiple comparisons; 120s: p=0.0470; 180s: p=0.0202; 240s: p=0.0016; 300s: p=0.0013; 360s: 0.0002; 420s: p=0.0003; 480s: p=0.0007; 540s: p=0.0020; 600s: p=0.0028; 660s: p=0.0094; 720s: p=0.0073; 780s: p=0.0155; 900s: p=0.0105][Extinction 2: Sidak’s multiple comparisons; 240s: p=0.0039; 300s: p=0.0187; 420s: p=0.0181][Extinction 3: Sidak’s multiple comparisons; 360s: p=0.0094])(Figure 5C, F, I). We again fit a GLM to determine if any variables could partially describe our calcium signals. We found that the best fit model included the isosbestic channel and kinematic information(R^2^_CS_=0.1536)(Fig 5K). Interestingly, freezing initiation and termination splines no longer significantly explained any variance within the model, suggesting the astrocytic signals were becoming decoupled to these moments. Through repeated exposures to the context, while calcium events are still present, the amount of information that is able to be decoded from them decreases, suggesting that these signals are more relevant to internal states. Overall, our data suggest that astrocytic calcium remains elevated even as freezing levels decline naturally across extinction sessions.

**Figure 4.**
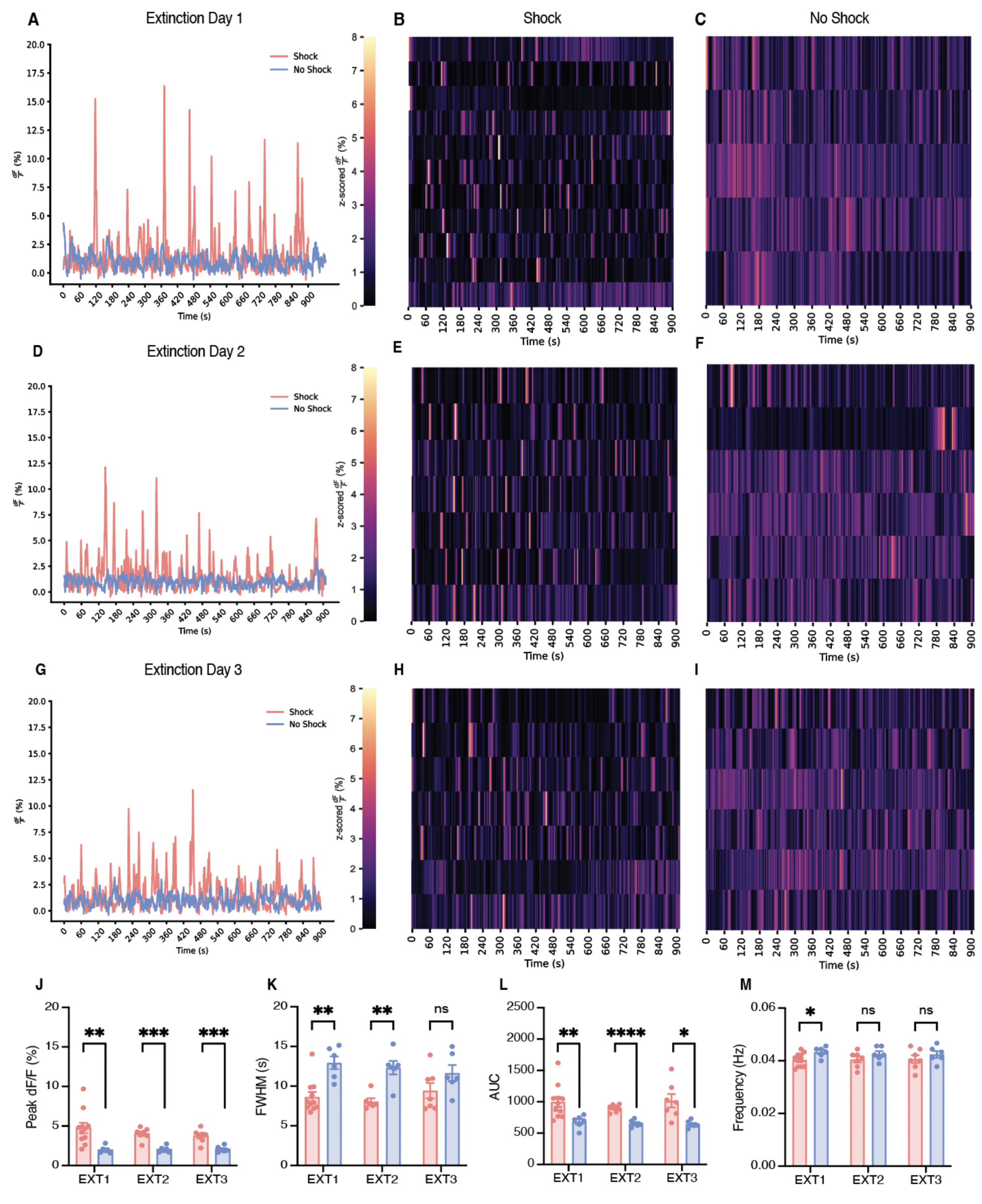
BLA astrocytes in the shock condition exhibit increased peak height, decreased duration, and increased total fluorescence of events compared to no-shock, but these do not change across extinction days. (A, D, G) Representative calcium time series (dF/F %) for shock and no-shock conditions during the 900 second contextual extinction sessions; (A) extinction day 1, (D) extinction day 2, (G) extinction day 3. (B, E, H) Z-scored dF/F (%) across extinction for shock condition; each row represents a single subject across time within the session. (C, F, I) Z-scored dF/F (%) across extinction for the no-shock condition; each row represents a single subject across time within the session. (J-M) Calcium event metrics; (J) peak height, (K) full-width half maximum, (L) area under the curve, and (M) frequency across all three days of extinction. Error bars indicate SEM. For t-tests, p ≤ 0.05, **p ≤ 0.01, ***p ≤ 0.001, ****p ≤ 0.0001, ns = not significant. *Extinction 1*: shock n=11, no-shock=6 (For the no-shock group, one animal’s recording was approx 90 seconds short. This animal was excluded from the raster plot and behavioral analysis, but still used for event metric calculations). *Extinction 2*: shock n=7, no-shock=6. *Extinction 3*: shock n=7, no-shock=6.

**Figure 5.**
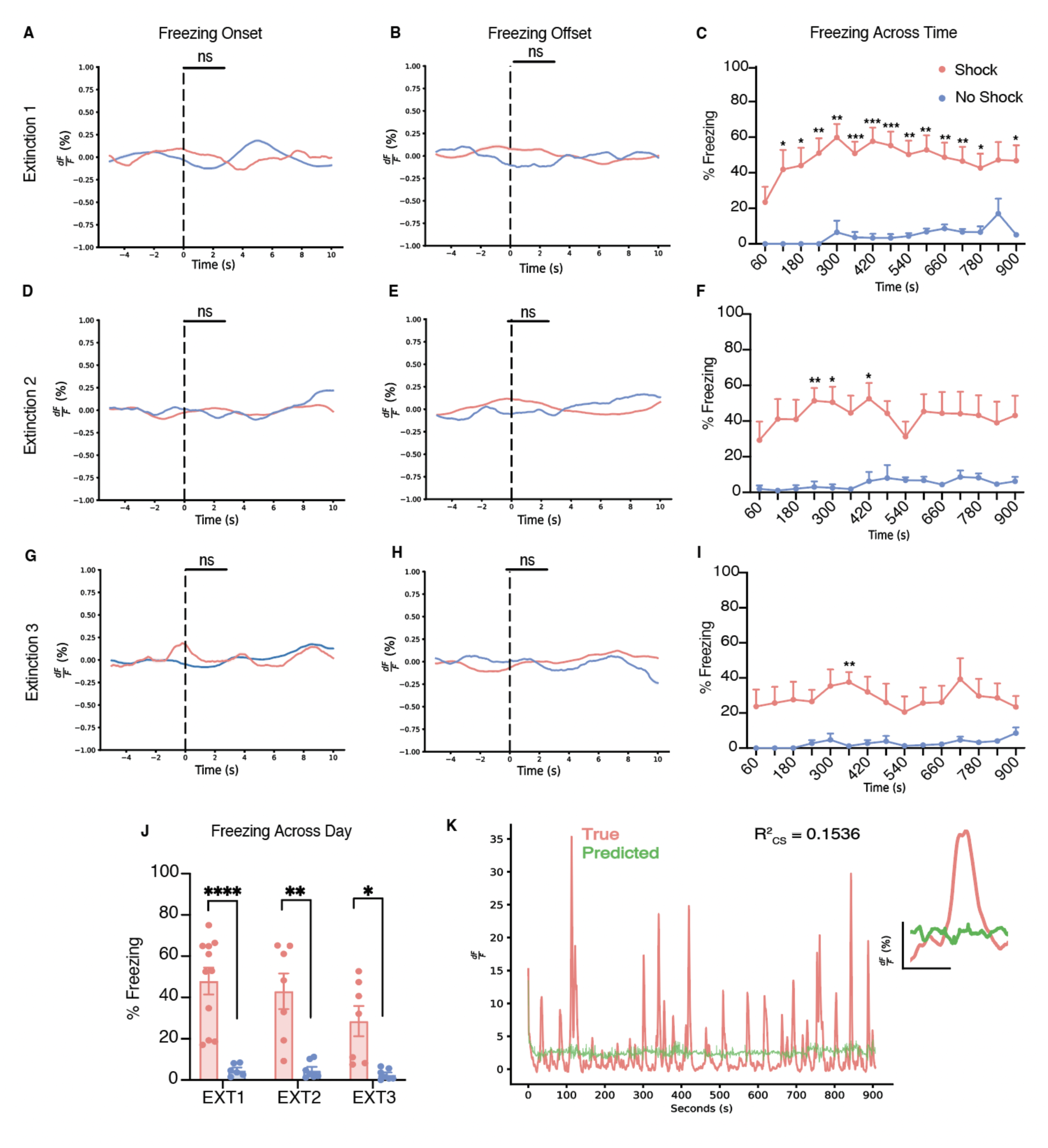
BLA astrocytic calcium does not respond to the initiation or termination of freezing behavior during extinction sessions. (A-B) Peri-event analysis for the initiation (A) and termination (B) of freezing behavior, with each event occurring at the dashed line (time = 0) for extinction day 1. (D-E) Peri-event analysis for the initiation (D) and termination (E) of freezing behavior, with each event occurring at the dashed line (time = 0) for extinction day 2. (G-H) Peri-event analysis for the initiation (G) and termination (H) of freezing behavior, with each event occurring at the dashed line (time = 0) for extinction day 3. (C, F, I) Percent freezing across time within (C) extinction day 1, (F) extinction day 2 and (I) extinction day 3. (J) Average percent freezing across three days of extinction for shock and no-shock conditions. (K) True and predicted traces produced from a generalized linear model. Error bars indicate SEM. For t-tests and ANOVAs, p ≤ 0.05, **p ≤ 0.01, ***p ≤ 0.001, ****p ≤ 0.0001, ns = not significant. For peri-event metrics, * = 95% CI; ** = 99% CI; *** = 99.9, ns = not significant. Extinction 1: shock n=11, no-shock=6 (For the no-shock group, one animal’s recording was approx 90 seconds, short. This animal was excluded from the raster plot and behavioral analysis, but still used for event metric calculations). Extinction 2: shock n=7, no-shock=6. Extinction 3: shock n=7, no-shock=6.

To further explore the memory-dependent nature of the observed astrocytic calcium changes, a subset of mice from the shock and no-shock groups were placed into a novel open field context (Cxt B) on Day 6 of our behavioral paradigm while recording astrocytic calcium (Figure 1H; Figure 6A-C). Here, we did not observe any significant differences in distance traveled, mean speed, number of center entries or total time spent in the center of the open field across shock and no-shock conditions ([Distance: Mann-Whitney U-test, p>0.9999][Mean speed: Mann-Whitney U-test, p>0.9999][Center entries: Independent t-test, p=0.9502][Center time: Mann-Whitney U-test, p>0.9999])(Figure 6D-G). Further, most mice in both groups did not freeze at all for the entirety of the open field session; thus we did not further analyze freezing behavior between groups or perform peri-event analysis with astrocytic calcium for freezing initiation or termination, as performed with other sessions. Most importantly, we assessed calcium event metrics during this session to observe if changes observed during contextual fear learning and extinction are due to an enhancement in activity maintained over time. Here, we did not observe any significant differences in the peak height, AUC, FWHM or frequency of calcium events across the shock and no-shock conditions ([Peak height: Mann-Whitney U-test, p=0.3429][FWHM: Independent t-test, p=0.6002][AUC: Independent t-test, p=0.5949][Frequency: Mann-Whitney U-test, p=0.9603])(Figure 6H-K). This suggests that astrocytic calcium activity changes due to shock are context-dependent and these cells are storing information related to the memory of the aversive experience.

**Figure 6.**
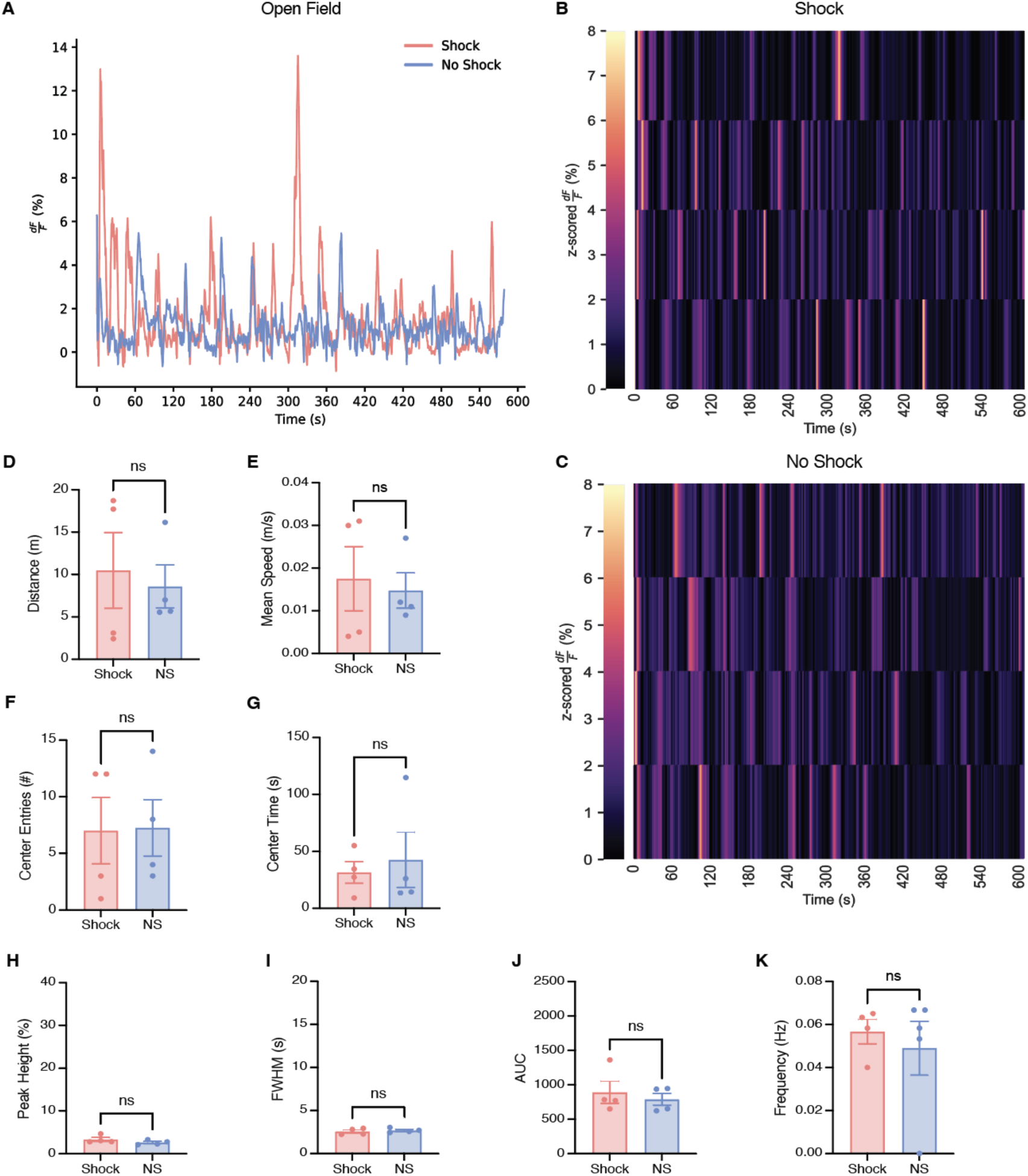
Astrocytic calcium event characteristics and behavior in a novel open field environment do not differ between shock and no-shock groups. (A) Representative calcium time series (dF/F %) for shock (coral) and no-shock (blue) conditions during the 600 second novel open field context B (Cxt B) session. (B-C) Z-scored dF/F (%) in Cxt B for shock and no-shock conditions; each row represents a single subject across time within the session. (D-G) Behavioral measures; (D) distance traveled (m), (E) mean speed (m/s), (F) number of center entries, and (G) time spent in the center. (H-K) Calcium event metrics; (H) peak height (%), (I) full-width half maximum (s), (J) area under the curve (AUC), and (K) frequency (Hz). (L-N) Peri-event analysis; (L) initiation, (M) termination of freezing behavior and (N) center entry. Error bars indicate SEM. For t-tests, p ≤ 0.05, **p ≤ 0.01, ***p ≤ 0.001, ****p ≤ 0.0001, ns = not significant. Open Field Cxt B: shock n=4, no-shock=4.

To further elucidate the role of astrocytes in context-dependent memory recall, we utilized the Tet-Tag activity-dependent labeling strategy to chemogenetically inhibit a fear memory while simultaneously performing astrocytic fiber photometry recordings (Figure 7A; left). This system relies on the coupling of the *cfos* promoter to the tetracycline transactivator (tTA), which in its protein form, binds directly to the tetracycline response element (TRE) in a doxycycline (Dox)-dependent manner and can drive expression of a protein of interest (e.g. hM4Di and/or mCherry)(Figure 7A; left). Moreover, the expression of the Designer Receptor Exclusively Activated by Designer Drugs (DREADDs), hM4Di, allows for chemogenetic silencing of the tagged experience (i.e., fear) during recall.

**Figure 7.**
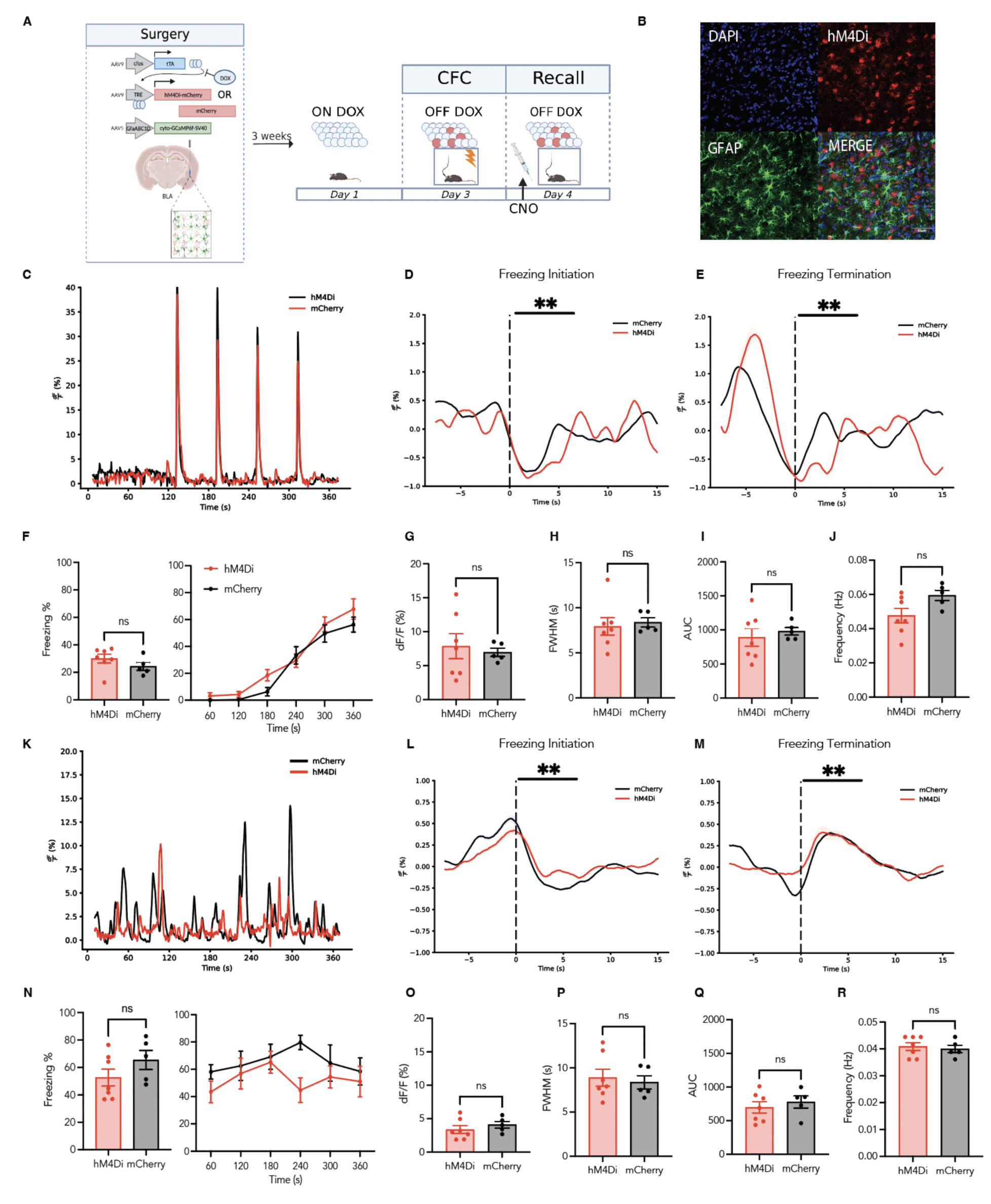
Fear engram inhibition in the basolateral amygdala (BLA) during recall does not modify freezing behavior or astrocytic calcium event characteristics. (A) Surgical and behavioral schematic; (Left) The genetically-encoded calcium indicator, AAV5-GfaABC1D-cyto-GCaMP6f-SV40, was unilaterally injected into the basolateral amygdala (BLA), in combination with bilateral injection of either AAV9-c-fos-tTA-TRE-hM4Di-mCherry (hM4Di) or AAV9-c-fos-tTA-TRE-mCherry (mCherry) control virus to allow chemogenetic control of labeled cells while recording astrocytic calcium dynamics. (Right) On Day 1, mice were taken off of their doxycycline (Dox) diet to allow for the opening of the labeling window 48 hours in advance of behavioral testing. On Day 3, mice underwent contextual fear conditioning (CFC) on Day 1 in Context A (Cxt A) for 360 seconds where they received 4, 1.5mA foot shocks. They were immediately placed back on their Dox diet, thus closing the labeling window. Day 4, mice were placed back into Cxt A for contextual recall for 360 seconds in the absence of foot shock. 30 minutes before this session, clozapine-N-oxide (CNO) was administered at 3 mg/kg to inhibit the labeled ‘engram’ during recall. 90 minutes after the start of the behavioral session, mice were perfused to capture peak endogenous cFos protein levels. (B) Representative images of hM4Di-mCherry/GFAP co-staining (red and green, respectively) and DAPI+ cells (blue); scale bar indicates 50 micrometers (um). (C) Representative calcium time series (dF/F %) for hM4Di (red) and mCherry (black) conditions during the 360 second contextual fear conditioning (CFC) session. (D-E) Peri-event analysis for the initiation (D) and termination (E) of freezing behavior during CFC, with each event occurring at the dashed line (time = 0). (F) Behavioral analysis for CFC; (left) average percent freezing and (right) freezing across time within the recall session. (G-J) Calcium event metrics for CFC, (G) peak height, (H) full-width half maximum, (I) area under the curve, and (J) frequency. (K) Representative calcium time series (dF/F %) for hM4Di (red) and mCherry (black) conditions during the 360 second recall session. (L-M) Peri-event analysis for the initiation (L) and termination (M) of freezing behavior during recall, with each event occurring at the dashed line (time = 0). (N) Behavioral analysis for recall; (left) average percent freezing and (right) freezing across time within the recall session. (O-R) Calcium event metrics for recall, (O) peak height, (P) full-width half maximum, (Q) area under the curve, and (R) frequency. All error bars and bands indicate SEM. For t-tests and ANOVAs, p ≤ 0.05, **p ≤ 0.01, ***p ≤ 0.001, ****p ≤ 0.0001, ns = not significant. For peri-event metrics, * = 95% CI; ** = 99% CI; ns = not significant. hM4Di = 7, mCherry = 5.

As shown in our experiments above, we began by performing simultaneous astrocytic calcium recordings with fiber photometry through the expression of AAV5-GfaABC1D-cyto-GCaMP6f in the unilateral BLA (Figure 7A; left). Prior literature has demonstrated that manipulation of the BLA with perturbation techniques impacts behavioral freezing levels typically associated with context-dependent recall (Han et. al., 2009; Gore et. al., 2015; Liu et. al., 2022). We hypothesized that bilateral inhibition of tagged BLA cells expressing hM4Di during recall would disrupt astrocytic calcium dynamics that we observed in our first experiment. On Day 1, mice had Dox diet removed to ‘open’ the engram tagging window 48 hours later. On Day 3, mice underwent a 360 second contextual fear conditioning (CFC) session with the administration of 4, 1.5mA foot shocks, as described in our previous experiments while astrocytic calcium dynamics were recorded in BLA (Figure 7A; right). Immediately after this CFC session, mice were placed back on their Dox diet to ‘close’ the tagging window (Figure 7A; right). On Day 4, mice underwent a 360 second contextual recall session in the absence of foot shocks. 30 minutes prior to this session, mice received intraperitoneal (I.P.) injection of clozapine-N-oxide (CNO) at 3 mg/kg to inhibit the ‘tagged’ fear engram during behavior (Figure 7A; right). Expression of hM4Di-mCherry in bilateral BLA with astrocytic GCaMP was confirmed in each animal prior to behavioral and calcium time series analysis (Figure 7B).

When comparing astrocytic calcium levels during CFC, both hM4Di and mCherry groups displayed robust increases at the initiation of each foot shock (120s, 180s, 240s, 300s), as previously described above (Figure 7C). This was expected, as no CNO was on board during the CFC session and we anticipated replication of our above findings. Additionally, in both groups there was time-locking to behavioral freezing initiation and termination epochs during CFC in both hM4Di and mCherry groups, further replicating our prior experiment (Freeze initiation and termination peri-event analysis; 99% CI)(Figure 7D-E). As expected, hM4Di and mCherry mice displayed the same freezing levels on average (Independent t-test; p=0.2395) and across time during CFC (Two-way ANOVA RM; Interaction: F (5, 50) = 1.107, p=0.3688; Time bin: F (2.623, 26,23) = 82.43, p<0.0001; Group: F (1, 10) = 1.564, p=0.2395; Subject: F(10,50) =3.685, p=0.0010)](Figure 7E). Finally, both groups displayed no significant differences in astrocyte calcium event characteristics (peak height, full-width half maximum, area under the curve, or frequency) during CFC, as expected ([Peak height: Independent t-test; Welch’s correction, p=0.6532][AUC: Independent t-test; Welch’s correction, p=0.5351][FWHM: Independent t-test; p=0.7292][Frequency: Independent t-test; p=0.0623)(Figure 8G-J).

For fear memory recall, mice in the hM4Di and mCherry groups did not display detectable changes in astrocytic calcium dynamics (Figure 7K). This was further confirmed by peri-event analysis at the initiation and termination of freezing bouts during recall. As shown in our above experiments, astrocytic calcium becomes time-locked to each of these behavioral epochs for both hM4Di and mCherry groups compared to their respective medians (Freezing initiation/termination: peri-event analysis, 99% CI)(Figure 7L-M). This suggests that CNO does not have an effect on astrocytic calcium dynamics during contextual recall. Interestingly, we did not observe any significant differences in behavioral freezing levels between the two groups, suggesting that chemogenetic inhibition of a fear engram in the BLA does not lead to a decrease in freezing levels (Independent t-test; p=0.2028), nor the recall of fear across time within session (Mixed-effects model (REML); Interaction: F (5,60) = 0.7030, p=0.6234; Time bin: F(1,60) = 5.257, p=0.0254; Group: F(5,60) = 0.6997, p=0.6258)(Figure 7N).

Furthermore, to quantify any differences in astrocytic calcium event metrics, peak height, FWHM, AUC and frequency (Hz) were analyzed. There were no significant differences in any of these metrics, with both groups mirroring the results described in our initial shock group findings ([Peak height: Independent t-test; p=0.3921][AUC: Independent t-test; p=0.5271][FWHM: Independent t-test; p=0.6981][Frequency: Independent t-test; p=0.6860])(Figure 7O-R). Overall, our data suggests that inhibition of a fear engram in BLA does not impact behavioral freezing levels, nor does it impact astrocytic calcium dynamics. It is important to note, however, that the freezing behavior and its tie to astrocytic calcium are maintained, strengthening our understanding of their potential role in behavioral expression of fear.

## 4) Discussion

Our results demonstrate that BLA astrocytes are differentially involved in the acquisition, recall, and extinction of a contextual fear memory. Strikingly, these astrocyte populations in the no-shock groups showed low levels of calcium dependent activity relative to the shocked group. These results corroborate previous research demonstrating that astrocytic populations are active specifically during salient experiences (Adamsky et. al., 2018). Furthermore, this could be related to an increased demand for neuronal metabolic support due to cellular activity recruited during memory formation. Recent models suggest that astrocytes become active during metabolically taxing experiences to support memory encoding and consolidation by providing astrocytically derived lactate to neuronal populations to increase ATP production (Steinman et. al., 2016; Adamsky & Goshen, 2018; Alberini et al. 2018). In line with this possibility, in our study mice that did not associate a noxious stimulus to the context displayed less activity than the shocked group. This could indicate that during memory acquisition, where a CS is paired with a US, astrocytic populations become involved to maintain this memory association. As the BLA preferentially parses salient information (Sengupta et. al., 2018; Pryce et. al., 2018), this could explain the lack of strong calcium events in the no-shock group, whereas in a structure that processes both associations and emotional salience (Zheng et. al., 2017; Eichenbaum, Schoenbaum, Young & Bunsey, 1996), such as the hippocampus, we predict to see reliable but increased events during contextual exploration and after CS-US pairing. Relatedly, and given that astrocyte assemblies are remarkably disengaged before the onset of foot shock during CFC in our study, future experiments may test whether populations of astrocytes are necessary for proper fear expression, and if after such salient experiences, BLA neurons require glial participation for stable memory correlates.

Interestingly, we only observed differences in the calcium event characteristics (e.g. FWHM, peak height) in the shock group when animals are undergoing CFC. This could indicate there are two distinct populations of cells, one which becomes active for memory encoding and maintenance, whereas the other is there to process incoming sensory input into the BLA or motor output. This hypothesis is supported by our modeling approach which reported significant contribution of kinematic related information as well as stimuli specific and relevant behavioral information. Furthermore, while the no-shock animals display minimal calcium transients these transients are significantly wider, possibly in response to exploring the novel context. This could be explained by the observation that AUC for these transients was also significantly lower for these animals, suggesting their was less overall calcium binding and subsequent recorded fluorescence, and thus indicating their may be cells that had sustained activity in response to a non-discrete stimuli, in contrast to something well defined like foot-shock. Furthermore, peri-events were observed in the fear conditioning and recall sessions. Specifically, we observe an increase in activity prior to the initiation and termination of freezing during recall. During recall, we observe an increase in activity prior to freezing initiation, but instead an increase in activity after the termination of freezing. It is possible that astrocytes ramp up their activity to either induce a state of freezing or become active in response to increased neuronal activation immediately prior to bouts of freezing. Regarding the termination of a freezing bout specifically, it is possible that astrocytes play a functional role in suppressing fear or anxiety states within the BLA (Cho et. al., 2022), though it is important to note that separate studies have also implicated their role in modulating locomotion (Qin et. al., 2020)--thus, as termination of freezing by definition requires movement, these two explanations remain to be reconciled. Notably, we did not observe these elevations in calcium activity at the initiation or termination of freezing during extinction. Indeed, we lose our predictive power in our GLM during extinction sessions, further evidencing this putative decoupling, and so it is likely these signals reflect internal physiological states that would be interesting to further investigate with higher spatial resolution. This could be due to the difference in session length causing changes to how these cells respond after the initial six-minutes, or it could be that after this initial exposure some subsequent learning has occurred causing these cells to respond to different local cues or internal states. While fiber photometry renders this possibility difficulty to test due to the lack of cell-specific granularity it affords, these hypotheses would be interesting to explore with higher resolution *in vivo* one-photon imaging approaches as it would allow us to visualize multiple subpopulations of BLA astrocytes, which could explain the diverse milieu of these signals.

Importantly, our findings that astrocytic calcium dynamics are modulated by salient stimuli were specific to the fear conditioned context, as supported by the lack of astrocytic calcium and behavioral changes in the neutral open field context. This is in line with previous findings showing that astrocytic manipulation enhances memory allocation in a task-specific manner during learning, but not in the home cage (Adamsky et. al., 2018). Astrocytic activity is heavily dependent on the presence of salient stimuli during a learning experience (i.e. foot shock) and that novel exploration of an environment is not sufficient to induce changes in the underlying calcium dynamics of astrocytes. This is congruent with data suggesting that the BLA shows sparse activity that is dependent on salient stimuli (i.e. reward) (Pratt & Mizumori, 1998).

Our findings suggest that chemogenetic-mediated engram inhibition in the BLA during recall does not abolish freezing behavior, nor affect astrocytic calcium dynamics. Future work may seem to express a pan-neuronal inhibitor throughout the BLA to test if this global reduction in activity modulates astrocyte dynamics, especially in light of data showing that BLA inhibition decreases freezing during recall. It is possible that the number of engram cells tagged in the BLA is not sufficient to abolish the behavioral expression of fear, or other brain regions may be compensating for or driving this behavior in coordination with the BLA. Astrocytes may still be intimately associated with engram cells in the BLA, but this perturbation is not sufficient to eliminate their activity. It would be interesting to label BLA engram cells and their associated astrocytes with chemogenetic tools to observe if the combination of cell inhibition (and thus, expanding the ‘engram’) abolishes memory recall, or if more temporarily precise optogenetic methods reveal an acute role of neurons in modulating astrocytes’ activity. Promisingly, such strategies that permit labeling and manipulation of both neurons and astrocytes will enable a better understanding of how heterogeneous cell types contribute to the overall memory engram as well as its behavioral expression.

While our results demonstrate a functional role of astrocytes in fear learning, the BLA is known to process additional salient information including fear, reward, novelty, etc. For instance, recent studies have shown that there are heterogeneous, genetically defined, populations within the BLA which may preferentially respond to a variety of stimuli and valences (Kim et. al., 2016). While our experiments did not tease out any valence-specific contributions of astrocytic calcium activity, future studies may deliver multiple valence-specific stimuli (e.g. sweetened condensed milk, social interaction, restraint stress) to animals while recording the corresponding calcium transients in the BLA. We posit that the BLA will display robust calcium dynamics to both positive and negative stimuli, albeit in partially separate populations of cells. This is consistent with recent literature showing that there are genetically-defined populations of cells along the anterior-posterior axis of the BLA that process fear and reward uniquely (Kim et. al., 2016).

While this work provides a role for astrocytes in conditioned fear, an ongoing issue surrounds whether astrocytes are mere support cells for neurons or if they actively encode information necessary for cognitive processes. Future experiments may concurrently record from neuronal populations to identify putative relationships between each cell type, and how real-time interplay between these populations supports learning and memory processes. A tantalizing possibility that combines neuron-glia relationships with neuromodulatory influences is that BLA astrocytes are necessary for proper adrenergic signaling which has been proposed in prior work (Gao et. al., 2016; Akther & Hirase, 2021). Also, it has been shown that CFC induces a downregulation of astrocytic Rac-1 (Liao et. al., 2017; Fan et. al., 2021), promoting astrocytic plasticity, which may explain our observed differences in calcium events between our two groups in that this increased astrocytic plasticity could be necessary for remodeling synaptic connections for continued signaling.

Finally, future studies may causally dissect the role of astrocytes by Gq or Gi pathway activation in these populations during recall or extinction to determine if cellular manipulation is capable of inducing either a memory enhancing or amnesic response to fear learning. As astrocytes do not have typical “inhibition/excitation” properties which are more typically associated with neurons (Durkee et. al., 2019; Van Den Herrewegan et. al., 2021), future research may take its amore complex signaling pathways into account and yield crucial information in how astrocytes participate at the tripartite synapse to facilitate the learning of conditioned fear. Indeed, higher resolution single-cell and populating imaging methods, combined with causal perturbation strategies, could be used to further delineate the role of these cells in memory formation and expression. Overall, our results suggest an active role of astrocytes in contextual fear learning within the BLA and reveal their dissociable role in contributing to memory recall and extinction.

## Acknowledgements.

This work was supported by a Ludwig Family Foundation grant, an NIH Early Independence Award (DP5 OD023106-01), an NIH Transformative R01 Award, a Young Investigator Grant from the Brain and Behavior Research Foundation, the McKnight Foundation Memory and Cognitive Disorders award, the Pew Scholars Program in the Biomedical Sciences, the Air Force Office of Scientific Research (FA9550-21-1-0310), the Center for Systems Neuroscience and Neurophotonics Center at Boston University.

## Data Availability Statement

The data that support the findings of this study are available from the corresponding author upon reasonable request.

## Code Accessibility

All statistical analysis and subsequent photometry analysis was performed via custom python scripts freely available at https://github.com/rsenne/RamiPho.

